# Outflanking Immunodominance to Target Subdominant Broadly Neutralizing Epitopes

**DOI:** 10.1101/346437

**Authors:** Davide Angeletti, Ivan Kosik, William T. Yewdell, Carolyn M. Boudreau, Vamsee V.A. Mallajosyula, Michael Chambers, Madhu Prabhakaran, Heather D. Hickman, Adrian B. McDermott, Galit Alter, Jayanta Chaudhuri, Jonathan W. Yewdell

**Affiliations:** Laboratory of Viral Diseases, National Institutes of Allergy and Infectious Diseases, National Institutes of Health, Bethesda, MD, USA; Immunology Program, Memorial Sloan Kettering Cancer Center, New York, NY, USA; Ragon Institute of MGH, MIT and Harvard, Cambridge, MA, USA; Harvard Ph.D. Program in Virology, Division of Medical Sciences, Harvard University, Boston, MA; Institute for Immunity, Transplantation and Infection, Stanford University, Stanford, CA, USA; Vaccine Research Center, National Institutes of Allergy and Infectious Diseases, National Institutes of Health, Bethesda, MD, USA; Laboratory of Clinical Immunology and Microbiology, National Institutes of Allergy and Infectious Diseases, National Institutes of Health, Bethesda, MD, USA

## Abstract

A major obstacle to vaccination to antigenically variable viruses is skewing of antibody responses to immunodominant epitopes. For influenza virus hemagglutinin (HA), the immunodominance of the variable head impairs responses to the highly conserved stem. Here, we show that head immunodominance depends on the physical attachment of head to stem. Stem immunogenicity is enhanced by immunizing with stem only-constructs or by increasing local HA concentration in the draining lymph node. Surprisingly, co-immunization of HA and stem alters stem-antibody class switching. Our findings delineate strategies for overcoming immunodominance with important implications for human vaccination.

---

Seasonal influenza remains a significant public health burden with vaccines requiring frequent reformulation yet providing limited protection (*1, 2*). Broadly neutralizing antibodies (Abs) binding the viral hemagglutinin (HA) have sparked the hope of developing a universal influenza vaccine (*3*). Most Abs target the highly variable globular head (*3–6*). The conserved HA stem is much more cross-reactive between strains but is poorly immunogenic following infection or vaccination (*7*).

HA and other immunogens activate naïve B cells present in lymph nodes (LNs) or spleen. Epitopes with sufficient avidity for B cell receptors (BCRs) trigger signaling events that lead to B cell seeding of germinal centers (GC). Here, B cells proliferate and experience somatic BCR hypermutation (SHM) and class-switch recombination (CSR). B cell clones with increased BCR avidity for immunogen are selected for proliferation and can differentiate into antibody-secreting plasma cells and memory B cells (*8*).

Although Abs can potentially bind to all surfaces of immunogenic proteins, Ab responses focus on a limited number of immunodominant antigenic sites. This phenomenon, termed immunodominance, is just now being defined and mechanistically dissected at the level of serum Abs and B cell responses (*9*). A recent study suggests that B cell precursor frequency and BCR avidity contribute to the subdominance of conserved HIV GP160 epitopes (*10*). Other studies suggest, however, that GC are more permissive than previously thought, allowing B cells with BCRs of even 100-fold differences in avidity to emerge from the same LN (*11, 12*).

To better understand the immunodominance of the HA head domain, we immunized mice intramuscularly (i.m.; the typical route for human vaccination) with full length PR8 H1 HA or native stem-only recombinant purified proteins (*13, 14*) in different combinations (Fig. 1A). 21 d after a single boost, we quantitated antigen-specific GC B cells in draining LNs, specific for HA head or HA stem via combinatorial flow cytometry staining using HA H1 and HA H5 (two proteins with different heads but semi-conserved stem) (Fig. S1A). GCs formed in the ipsilateral draining LNs in similar numbers independently of the immunogen (Fig. S1B-C). As expected (*5, 15, 16*), HA i.m. immunization exclusively induced head-specific GC B cells. Importantly, stem immunization generated a stem-specific B cell response of similar magnitude to the head specific response, demonstrating that our stem construct is not intrinsically of low immunogenicity (Fig. 1B). Consistent with a previous study (*12*), we could not account for the specificity of a large number of GC B cells, which may recognize non-native forms of the immunogens, self/microbiome derived antigens, or immunogen contaminants (Fig. 1B). After immunization with HA and stem in separate legs, immunogen-specific B cells developed only in the ipsilateral LN (Fig. 1C and Fig. S2A-B). Notably, this lack of competition in the draining LN also occurred when we mixed intact HA and stem, (Fig. 1C and Fig. S2A-B), strongly suggesting that that head immunodominance results from naïve B cell competition for full-length HA.

**Figure 1.**
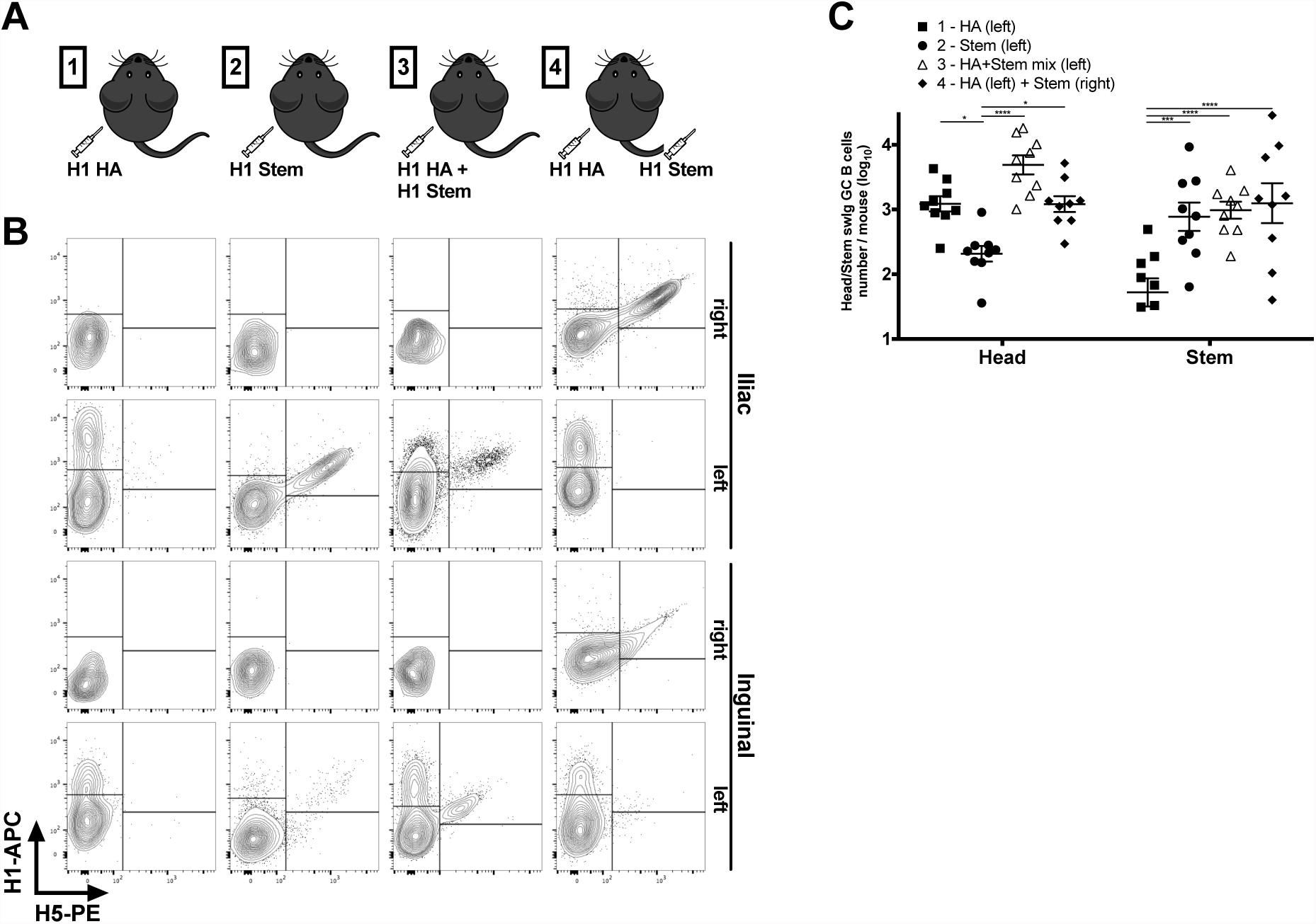
Immunodominance of the B cell responses depends on physical attachment of the HA head to stem. (A) Schematic of the immunization strategy. Group 1 was immunized with full length HA in the left hind leg, group 2 with stem in the left hind leg, group 3 with full length HA and stem in equimolar amount in the left hind leg and group 4 with full length HA in the left hind leg and stem in the right hind leg. (B) Representative flow cytometry plot showing swig GC B cells, gated as live CD3^-^ B220^+^ GL7^+^ CD38^-^ IgD^-^ IgM^-^, ability to bind HA head (H1 single positive) and HA stem (H1^+^H5^+^). (C) Enumeration of head vs stem swig GC B cells for the 4 different groups 21 days after challenge. Three independent experiments with 4 mice each (pooled for the first experiment) (n=9). Bar is mean ± SEM, statistical analysis was performed using two-way ANOVA with Holm-Sidak multiple comparison test.

We next measured head *vs*. stem specific serum Abs in the 4 immunization groups (Fig. 2A). Stem was highly immunogenic as a standalone immunogen and HA failed to elicit a stem response in 11 of 12 mice. Stem-induced Abs bound to the H5 HA (Fig. S3A) and inhibited binding of two well characterized anti-stem mAbs (Fig. S3B) (*17, 18*). Ab titers partially correlated with GC B cell numbers (Fig. S3C) suggesting bottlenecks between GC B cells, PC differentiation and serum Ab production. Interestingly, mixing HA with stem revealed a further discrepancy between GC B cell numbers and serum Ab titers, as stem specific titers were reduced 10-fold relative to stem only immunization. This suggests possible competition for help from T follicular cells (*19*) and follicular dendritic cells (*20*) essential for PC differentiation, or competition at post-GC step in serum Ab secretion.

Since anti-stem Ab-based protection in mice is mediated by Fc-mediated functions (*21*), Ab heavy chain class is critical in anti-stem Ab mediated protection. HA predominantly induces mouse IgG1 and IgG2 Abs, with only the latter able to interact with Fc receptors and activate complement. Remarkably, we found that head-specific Abs induced by HA immunization were biased to IgG1 while stem-induced Abs were IgG2 biased, a pattern that was maintained when immunizing with each immunogen in opposite legs (Fig. 2B). Importantly, however, stem Abs induced by HA/stem mixed immunization shifted towards IgG1, providing an important potential caveat for mixing immunogens in a single site.

**Figure 2.**
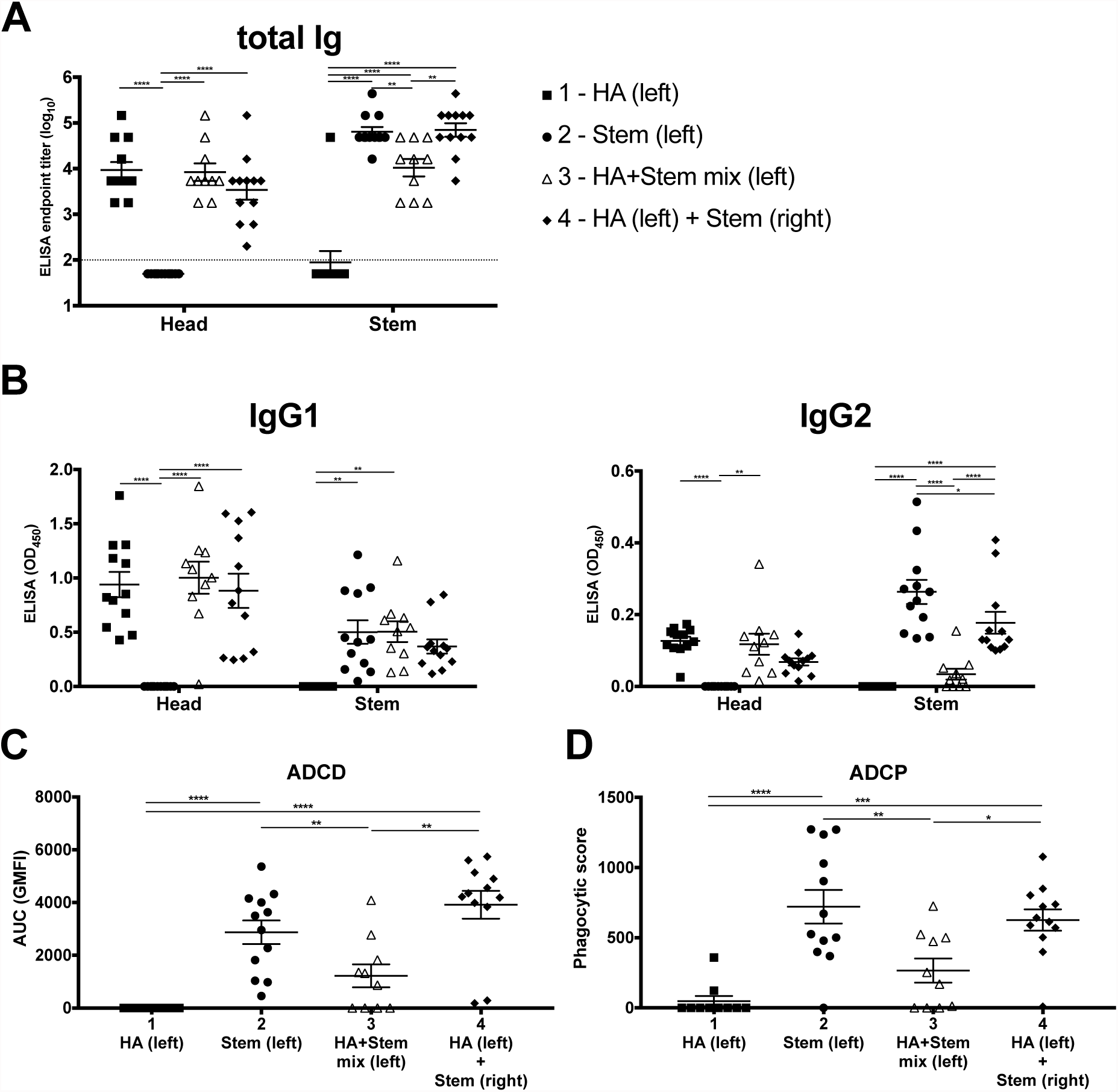
Serum immune responses of immunized mice is impaired upon mixed immunization. Antibody endpoint titers (A) and IgG subclass response (B) to HA head (following stem Abs absorption) and to stem for the different immunization groups (n=12 for groups 1, 2, 4 and n=10 for group 3). Bar is mean ± SEM, statistical analysis was performed using two-way ANOVA with Holm-Sidak multiple comparison test. (C) Sera were tested for the ability to induce ADCD on stem-conjugated beads. Data is presented as area under the curve (AUC) of geometrical mean fluorescent intensity (GMFI) of 1:5 and 1:10 dilutions and each data point is the mean of two technical replicates (D) Ability of the sera to induce ADCP on stem-conjugated beads by primary monocytes. Each data point is the mean of two technical replicates. Three independent experiments with 4 mice each (n=12 for groups 1, 2, 4 and n=10 for group 3). Bar is mean ± SEM, statistical analysis was performed using one-way ANOVA with Tukey multiple comparison test.

Stem-induced sera were unable to neutralize PR8 *in vitro* (Fig. S4A) (*13, 17*), despite binding to HA on cell surface (Fig. S4B). Rather, *in vitro* serum neutralization titers highly correlated with head-specific Ab ELISA titers (Fig. S4C). We tested sera for Ab dependent complement deposition (ADCD) activity and Ab dependent cellular phagocytosis (ADCP) by primary monocytes (*22*). We found that stem-only immunization induced high levels of stem-dependent ADCD and ADCP and that mixed immunization significantly reduced stem-mediated effector functions (Fig. 2C-D). This revealed a strong correlation with IgG2 anti-stem titers, but not total IgG titers (Fig. S5A-B). When sera were tested on H5-HA-conjugated beads, ADCD activity mirrored the stem results (Fig. S5C), however PR8-HA ADCD and ADCP results highlighted the important contribution of anti-head Abs (Fig. S5D-E).

Taken together, these findings indicate that mixing immunogens at the same immunization site or immunizing at distant sites can influence the magnitude and quality of Ab responses.

Why does the physical attachment of the head to the stem suppress Ab responses to the stem? We hypothesized that head immunodominance results from sequestration of full-length HA by naïve head-specific B cells, preventing stem-specific B cell activation due to differences in naïve B cell numbers or BCR avidity (*9*). We first measured the frequencies of head- and stem- specific naïve B cells. Using a method developed for quantitating naïve T cells (*23*), we detected naïve HA-specific B cells at low frequencies, despite their low avidity. B cells specific for HA were detected by double staining with the same probe labelled separately with different fluorophores (Fig. S6A), thus excluding B cells specific for streptavidin and fluorophores. This revealed that HA and stem precursor frequencies are similar: ~250/300 cells per million naïve mature B cells (Fig. S6B-C).

We next measured GC B cell avidity at day 10 after full-length HA or stem immunization using the AC_50_ method (*24*). This was the first day post-immunization with sufficient expansion of GC B cells to identify antigen-specific class-switched (swIg) and IgM cells in draining LN (Fig. S7A). Consistent with our prior findings (*24*), head-specific B cells exhibited AC_50_ values in the low nM range. Critically, the avidity of stem-specific B cells was ~ 10-fold lower (Fig. S7B-C). Consistent with their higher avidity, the median fluorescent intensity (MFI) of HA^+^ cells was significantly higher for head-specific B cells (Fig. S7F). By 3 weeks post- boosting, the avidity of both head- and stem- specific GC B cells had increased 5-20-fold, as expected from affinity maturation (Fig. S7D-E).

Our results partially contradict recent reports that responding GC B cells can secrete mAbs with 10- to 100-fold differences in avidity for immunogen (*11, 12*). However, there are several possible explanations for this difference: firstly, while we measure BCR avidity other studies use soluble mAb, therefore BCR multimerization might contribute to the higher affinity we measure here. Secondly, by immunizing with stem only, we allow stem B cells to affinity mature without competition, thus reaching a higher affinity. Finally, a study, where relevant naïve B cell affinities were measured, shows that only higher affinity precursors can enter GC for monomeric antigen (as the ones used here) and, even for multimeric antigen, a 30-fold naïve avidity difference was sufficient to exclude low avidity B cells from GC (*10*).

Importantly, the contradictory findings above (*11, 12*) derived from footpad (f.p.) immunization, while we used i.m. immunization. To determine how the route of immunization influences antigen concentration in the draining LN, we immunized mice i.m. or f.p. with 10 µg R-phycoerythrin (PE) in adjuvant and 24 h later measured the amount of fluorescent PE present in LN extracts (Fig. 3A). Strikingly, after f.p. immunization we could detect PE in 78% of the popliteal LNs and 61% of the iliac LNs tested, while after i.m. immunization PE was detectable only in 22% of the LNs analyzed (for both the inguinal and iliac) (*p*=0.0022; Fisher’s exact test). If we average the amount of PE detected (with our limit of detection being 0.1 ng/ml) in the popliteal LN after f.p. immunization was at least 14- and 30-fold higher, respectively, than in draining inguinal and iliac LN after i.m. immunization.

**Figure 3.**
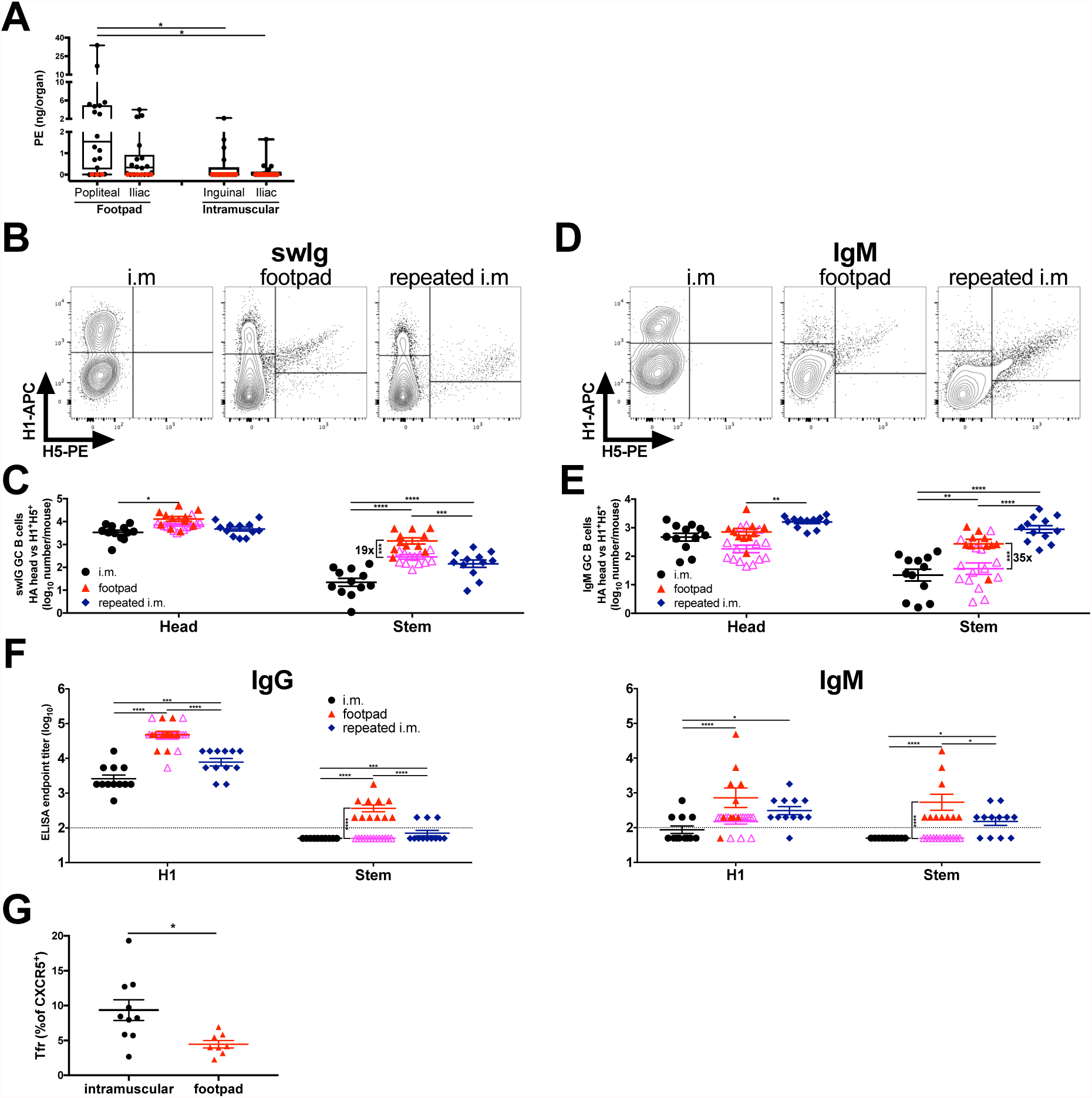
Stem subdominance can be subverted by increasing local antigen concentration. (A) amount of PE detected in draining LNs following f.p. or i.m. immunization (n=17-18). Box plot showing median and min to max. Red dots represent a value below the limit of detection which is indicated as zero. Statistical analysis was performed using one-way ANOVA with Tukey multiple comparison test. Representative flow cytometry plot showing swig GC B cells (gated as live CD3^-^ B220^+^ GL7^+^ CD38^-^ IgD^-^ IgM^-^)(B) and IgM GC B cells (gated as live CD3^-^ B220^+^ GL7^+^ CD38^-^ IgD^-^ IgM^+^)(D), ability to bind HA head (H1 single positive) and HA stem (H1^+^H5^+^) after i.m., footpad or repeated i.m. immunization. Enumeration of head vs stem swig GC B cells (C) and IgM GC B cells (E) for the 3 different groups 21 days after immunization (n=12 for i.m. and repeated i.m. and n=24 for f.p.). (F) Antibody endpoint titers for total Ig, IgG and IgM to H1-PR8 or stem for the different immunization groups. For f.p. red symbols indicate mice that are stem-seropositive while magenta indicates stem-seronegative (“low”). Three independent experiments with 4 mice each for i.m and repeated i.m and 5 independent experiments with 5 or 4 mice each for f.p. Bar is mean ± SEM, statistical analysis was performed using two-way ANOVA with Holm-Sidak multiple comparison test. (G) Quantification of Tfr contribution to total Tf (CD4^+^ CXCR5^+^ PD1^+^) (Tfr/Tfh ratio) in draining LN (iliac for i.m. and popliteal for f.p.) 8 days post immunization. (n= 10 for i.m. and 7 for f.p.). Two independent experiments with 5 mice each for i.m. and with 4 and 3 mice for f.p. Bar is mean ± SEM, statistical analysis was performed using unpaired t-test

We next immunized mice either f.p. with 10 µg of HA or i.m. with 10 µg of HA or i.m. with 25 µg of HA followed by and additional 50 µg of HA on day 1 and on day 2 to increase HA concentration in the LN (repeated i.m.). Three weeks post-immunization, GC frequencies in the iliac LN were similar but nodal cellularity significantly expanded with increasing Ag dose, with the average number of GC B cells per LN compared to i.m. immunization nearly doubling, or tripling, respectively, with f.p. or repeated i.m. immunization (Fig. S8A-C). Interestingly, while IgM^+^ GC B cells were almost absent after i.m. immunization, they were present in low numbers after f.p. immunization, and after repeated i.m. immunization increased to nearly equal switched Ig (swIg) after repeated i.m immunization (Fig. S8B). This suggests both with sustained recruitment of B cells to the GC due to increased immunogen delivery and with extended cycling of IgM^+^ GC B cells that are not able to receive adequate signals from Tfh to exit the GC reaction.

All immunization groups generated large numbers of head-specific IgG^+^ and IgM^+^ GC B cells on d 21, correlating with IgG- and IgM- head specific serum responses. In contrast, i.m. immunization generated low number of stem specific IgG and IgM GC B cells and undetectable stem-specific serum responses. F.p. immunization increased stem-specific switched B cells 100-fold compared to i.m. (Fig. 3B-C and Fig. S8D-E), comparable to stem only immunization (Fig. 1). Repeated i.m. immunization also increased switched, and IgM^+^ stem-specific stem B cells (Fig. 3D-E).

We were unable to detect IgG or IgM anti-stem Abs in i.m. immunized mice (Fig. 3F). f.p. immunization generated stem specific IgG and IgM serum responses in the ~50% of mice with the highest levels of GC B cells (on average 20x more abundant than seronegative mice (Fig. 3C)). This is consistent with a functional threshold of GC responses to generate sufficient numbers of plasmablast/plasma cells for a detectable serum Ab response. Based on cell surface phenotype by flow-cytometry, head- and stem-specific GC B cells had a similar dark zone (DZ) / light zone (LZ) distribution (Fig. S8F-G).

Consistent with the B cell analyses, repeated i.m. immunization elicited significant levels of H5 cross-reactive IgG in 3/12 mice; although 8/12 mice generated IgM responses. Although stem titers only partially correlated with H5 titers (Fig. S9A), all H5 reactivity was removed by competition with anti-stem mAbs. Immunization route also shaped the GC B cell immunodominance hierarchy among the 5 major head antigenic sites (Fig. S9B-C). At most extreme, Ca1 specific-responses nearly double in f.p. vs. i.m. immunization, consistent with our previous findings of route dependence of immunodominance (*16*).

Finally, we determined the influence of f.p. vs i.m. HA immunization on the ratio of T follicular regulatory cells (Tfr) to T follicular helper (Tfh) cells, which has been reported to regulate Ab responses (*25*). Although f.p. immunization generated higher frequencies of both Tfh and Tfr compared to i.m. (Fig. S10A-B-C), the Tfr/Tfh ratio was doubled after i.m. immunization, suggesting an additional contribution to the suppression of stem-specific response (Fig. 3G).

These findings demonstrate that the route of immunization governs the immunodominance of both the head vs. the stem regions. Stem subdominance is driven by differential B cell affinity under conditions of limiting immunogen. To circumvent immunodomination, immunizing with a subdominant protein is a promising approach for universal vaccination. While we can’t completely exclude that the lack of glycosylation of our bacterially synthesized stem construct accounts for its improved immunogenicity *vs.* stem in full length HA, this seems unlikely based on studies showing that, in the context of HA, stem immunogenicity is enhanced by blocking head epitopes with glycans (*26*) and not affected by removing stem glycans (*27*). Although mixing stem with HA does not interfere with head-specific responses, it reduces stem-specific polyclonal titers 10-fold and interferes with class switching, decreasing anti-stem ADCD and ADCP. Diminished immunogenicity from mixing of viral (*28, 29*) or protein vaccines (*30*) or immunization in different sites (*31*) clearly occurs in humans, but had not previously been mechanistically dissected.

In mice we could bypass affinity-based head immunodominance by immunizing in small anatomic space (the footpad) draining to a limited lymphshed. In humans, practical realities would dictate either single immunization with a subdominant immunogen, immunization in in different sites (legs and arms, for example) or engineering subdominant immunogens to increase LN delivery, either by modifying delivery (e.g. needle patches or slow release vaccines) or attaching immunogens to molecules/particles that enhance delivery to B cell areas of draining LN (*32*).

## Acknowledgements

We thank the NIAID Comparative Medicine Branch for maintaining the mice used in this study and Raghavan Varadarajan (Molecular Biophysics Unit, Indian Institute of Science, India) for assistance with stem protein design and production. This work is supported by the Division of Intramural research, National Institute of Allergy and Infectious Diseases

## Supplementary Information

### Experimental procedures

#### Animals

C57BL/6 mice were purchased from Taconic Farm. For all experiments female 8-12 weeks old mice were used and randomly assigned to experimental group. All mice were held under specific pathogen-free conditions. All animal procedures were approved and performed in accordance with the NIAID Animal Care and Use Committee Guidelines.

#### Proteins and immunization

Recombinant A/Puerto Rico/8/34 (PR8) HA with the Y98F mutation(*1*) and H1HA10-Foldon stem only construct derived from PR8(*2*) were used for the initial immunization studies. Animals were randomly divided in 4 groups and immunized i.m. in the left hind leg, under mild isoflurane anesthesia, with recombinant proteins mixed with Titermax Gold adjuvant (Sigma). Group 1 and 4 received 10ug of HA (0.8uM) in the left hind leg, group 2 received 4ug of stem (0.8uM) in the left hind leg while group 4 the same amount in the right hind leg. Group 3 received equimolar mixture of the two proteins in the left hind leg. 21 days after prime all the groups were boosted with the same amount of protein without adjuvant. 21 days after boost, serum was collected, animals sacrificed and spleen and inguinal and iliac LN collected.

Other groups of animal were immunized with HA sucrose purified from PR8 virus(*3*). After addition of PBS the solution was spun at 50,000x*g* for 2h at 4 C. The supernatant, containing HA and NA, was incubated at 56 C for 30 min after addition of 10mM EDTA. Purity was verified by Western blot and HA quantified by DC Protein Assay (Biorad). Animals were then immunized i.m. or via the hock f.p. with 10ug of HA (0.8uM) mixed with Titermax Gold adjuvant (Sigma). For repeated i.m. animals were immunized i.m. with 25ug of HA with Titermax Gold adjuvant on day 0, followed by 50ug of HA without adjuvant on day 1 and 50ug of HA without adjuvant on day 2. On day 21 serum was collected, animals sacrificed and spleen and iliac LN collected with also inguinal for i.m. and popliteal for f.p.

#### Flow cytometric analysis

For flow cytometric detection recombinant PR8 H1 HA and A/Indonesia/5/05 H5 HA with Y98F mutation(*1*) were pre-labelled with streptavidin-APC and streptavidin-PE (ThermoFisher, SA1005 and SA10041), respectively. Briefly, proteins were diluted at 0.02mg/ml and labelled by stepwise addition (at 10 minutes intervals) of molar excess of fluorescent streptavidin. Labelled proteins were stored at 4°C and generally used within 2 weeks of labelling.

Draining LN and spleens were harvested individually and single cells suspension prepared by mechanical dissociation. Red blood cells were lysed using ACK lysis buffer (Gibco). Cells were counted and up to 5 million cells stained per tube. Organs were stained with fluorescent HA (final concentration 4nM for each probe) and Ab cocktail for 1h at 4°C in PBS/0.1% BSA. The following Abs were used in different experiments: from BD Bioscience BV510 anti mouse CD3 (clone 145-2C11)(cat 563024), APC-Cy7 anti-mouse B220 (cloneRA3-6B2)(cat 552094), Pacific Blue anti-mouse B220 (cloneRA3-6B2)(cat 558108), FITC anti-mouse CD38 (clone 90)(cat 558813), FITC anti-mouse GL7 (clone GL7)(cat 553666), PE anti-mouse GL7 (clone GL7)(cat 561530), PerCP-Cy5.5 anti-mouse IgD (clone11-26c.2a)(cat 564273); from Biolegend BV785 anti-mouse B220 (cloneRA3-6B2)(cat 103246), Pacific Blue anti-mouse IgD (clone11-26c.2a)(cat 405712), APC/Fire750 anti-mouse IgD (clone11-26c.2a)(cat 405744), PE/Dazzle594 anti-mouse IgD (clone11-26c.2a)(cat 405742), PE/Dazzle594 anti-mouse IgM (clone RMM-1)(cat 406530), BV606 anti-mouse IgG (poly4053)(cat405327), Pe-Cy7 anti-mouse CD86 (clone GL-1)(cat 405014), PE anti-mouse PD-1 (clone 29F.1A12)(cat 135205), FITC anti-ICOS (clone C398.4A)(cat 313506); from eBioscience PerCP-e710 anti-mouse CD38 (clone 90)(cat 46-0381-82), e450 anti-mouse GL7 (clone GL7)(cat 48-5902-82), PE-Cy7 anti-mouse IgM (clone II/41)(cat 25-5790-82), e450 anti-mouse CXCR4 (clone 2B11)(cat 48-9991-82), APC-e780 anti-mouse CD4 (clone GK1.5)(cat 47-0041-82). After PBS washes, live/dead staining was performed with Live/Dead fixable Aqua kit (Thermo Fisher cat L34957) for 30 minutes at 4°C. Cells were washed trice with PBS/0.1% BSA and resuspended in PBS/0.1% BSA. Samples were analyzed using a BD LSR Fortessa X-20 instrument. Analysis was performed using FlowJo software (TreeStar).

For staining of T follicular cells, after single cell suspension preparation, cells were stained with antibody cocktail in 25ul + Biotin anti-mouse CXCR5 (clone SPRCL5)(eBioscience cat 13-7185-82) at 1:25 dilution in PBS/0.1% BSA. After washes, cells were incubated with streptavidin-APC for 15 min at 4°C, washed and live/dead staining performed as above. Cells were then permeabilized with eBioscience Foxp3 staining buffer set (cat 00-5523-00) according to manufacturer’s protocol. After permeabilization cells were washed with Permeabilization Buffer and stained with PE-Cy7 anti-mouse Foxp3 (clone FJK-16s)(eBioscience cat 25-5773-82) at 1:25 dilution for 30 min at RT. Cells were finally washed with permeabilization buffer and resuspended in PBS/0.1% BSA for analysis as above.

#### Precursor frequency quantification

For precursor frequency quantification we took advantage of the method developed for TCR (*4*) in order to increase staining of otherwise low affinity cells. Spleen, inguinal, axillary, mandibular, cervical and mesenteric LNs were collected from naïve mice, pooled and prepared as above. Protein kinase inhibitor (PKI) dasatinib (Axon Medchem) was added at a final concentration of 50 nM for 30 min at 37°C. Without washing, 10ug of unlabeled Streptavidin (Invitrogen, cat 434301) were added per tube for 20 min at 4°C. Again, without washing, proteins were added in the following combinations with final HA concentration of 8nM for 40 min at 4°C: H1-APC + H1-PE, H5-APC + H5-PE, H1-APC + H5-PE (to determine stem precursor frequencies), SA-APC + SA-PE (to determine SA precursor frequency), H1-APC + SA-PE (control). Cells were washed twice and unlabeled anti-mouse-PE (clone PE001)(Biolegend cat 408101) and anti-mouse APC (clone APC003)(Biolegend cat 408001) were added at 10ug/ml for 20 min at 4°C. After washes cells were stained with the following surface Abs: from Biolegend PE/Dazzle594 anti-mouse CD3 (clone 17A2)(cat 100246), FITC anti-mouse CD21/CD35 (clone 7E9)(cat 123407), APC/Cy7 anti-mouse CD23 (cloneB3B4)(cat 101630); from BD Bioscience BV421 anti-mouse CD43 (clone S7)(cat 562958); BV785 anti-mouse B220, PE-Cy7 anti-mouse IgM, PerCP-Cy5.5 anti-mouse IgD (as above). Aqua live dead staining was performed as above and more than 10 milion cells acquired per tube. Precursor frequency was defined as PE/APC double positive population – the frequency of the same population in the control tube.

#### AC_50_ measurement of GC B cell population affinity

Draining LN were prepared and stained as above, using a low input cell number per tube (10,000-20,000 antigen specific cells per tube) and incubated with a graded concentration of unlabeled rHA (0.66nM to 66nM). After washes, HA was detected using APC-conjugated streptavidin at 1:500 dilution for 30 min at 4°C. Data were plotted using frequency of rHA positive B cells and 50% maximal binding (AC_50_) calculated using a single one-site binding with Hill slope calculation (*5*).

#### ELISA and serum quantification

Microlon medium binding half-well ELISA plates (Greiner Biotech) were coated overnight at 4°C with recombinant H1, H5 HAs and stem (*6*) in 25ul PBS. Plates were blocked with 50ul PBS/4% milk for 2 hours at RT. After 3x washes with PBS+0.05% Tween-20 (PBST) plates were incubated with three-fold dilutions of sera starting from 1:200 in PBST for 90 min at RT. After 3x washes plates were incubated with 25ul of rat anti-mouse kappa, IgG1, IgG2(b+c) or IgM specific HRP-conjugated (Southern Biotech) or peroxidase anti-mouse IgG (vectorlab) diluted 1:2000 for 1h at RT. After 3x washes plates were developed for 5 minutes using TMB substrate (KPL biomedical) and halted with 0.1N HCl. Plates were read at 450nm. For endpoint titer determination, sera from at least six mice immunized with irrelevant proteins were tested using the same conditions. Cutoff for positivity was determined using the formula from (*7*). For quantification of head Ab titer, the serum was pre-absorbed using recombinant stem as follows. 5ug of his-tagged stem was incubated with Talon-Agarose resin in PBS, following 1h incubation at 4°C the slurry was washed with PBS and then incubated with sera o.n. in rotation at 4°C. The resin was spun at 300xg and allowed to settle and the supernatant collected and used for ELISA. For each sera a pre- and a post- absorption samples were run on all the recombinant proteins to verify depletion. For endpoint titer determination “Head titer” was considered based on reactivity of depleted sera on H1 while “stem titer” on the reactivity of serum pre-depletion on stem.

For competition with stem mAbs FI6 and 310-16G8 ELISA was performed as above with the following modifications: H1 HA coated plates were incubated with pooled HA (group 1) or stem (group 2) sera, after washed the human stem mAbs were dispensed to the wells at the concentration giving 75% of their maximum signal. Plates were detected with anti-human-kappa (southern biotech) diluted 1:100 and results express as % binding relative to uncompleted mAb.

#### ELISA Immunodominance profile

Plates were coated with equal amount of virally purified HA derived from D4 viruses as described in (*8*). After blocking, plates were incubated with two-fold serially diluted sera starting from 1:100 for 1h at RT. Following washes with PBST, serum binding was detected using anti-mouse-kappa secondary Ab at 1:2000 for 1h at RT. ELISA binding is expressed as area under the curve, calculated using GraphPad Prism as described before (*8*).

#### Antibody Dependent Cellular Phagocytosis (ADCP) assay

Biotinylated proteins were incubated with 1mm fluorescent neutravidin-coated beads (Invitrogen cat no F8776) for 2 hours at 37°C. Antigen-coated beads were then incubated with serum samples diluted 1:50 in PBS for 3 hours at 37°C in 96-well plates. Unbound antibody was washed away, and primary BALB/C mouse monocytes (isolated using EasySep Mouse Monocyte Isolation Kit, Stemcell Technologies cat no 19861) added at 12,500 per well and incubated at 37°C for 4 hours, then fixed. Phagocytosis was measured by flow cytometry on an Intellicyte iQue cytometer. Phagocytic scores are the % of bead positive cells x GMFI/10,000.

#### Antibody Dependent Complement Deposition (ADCD) assay

Antigen-coated beads were prepared as in ADCP and incubated with diluted serum samples for 2 hours at 37°C. Lyophilized guinea pig complement (Cedarlane) was resuspended in ice cold water, then diluted in veronal buffer with 0.1% gelatin, calcium and magnesium (Boston BioProducts cat no IBB-300X). Complement was added to opsonized beads and incubated for 20 minutes at 37°C. Beads were then washed with 15mM EDTA, stained with anti-guinea pig C3 (MP Biomedicals cat no 0855385), and incubated 15 minutes at room temperature. Samples were washed and analyzed on a BD LSRII cytometer with a high-throughput sampler to record the geometric mean of fluorescence intensity. Area under the curve was calculated using Graphpad Prism based on two dilutions (1:5 and 1:10)

#### PE quantification in LN

R-Phycoerythrin (Prozyme, cat PB32) was diluted in PBS and injected i.m. or via the f.p. 24h after injection, mice were sacrificed and draining LN removed and homogenized. Single cell LN suspension was filtered and resuspended in 200ul and 50 ul transferred to the well in two duplicate two-fold dilutions. A standard curve with 24 two-fold dilutions of R-PE starting from 1ug/ml was performed on every plate (Costar, White 96 well plates). Plates were read using Synergy H1 at ex 535nm / em 575 nm.

#### Virus inhibition assay

The self-reporting PR8-mCherry expressing virus was employed to measure neutralization properties of immune sera. Briefly, 40,000 MDCK cells were seeded on black 96 well plate. The mCherry expressing virus (500 TCID50) was preincubated with serial dilutions of sera at 37°C for 60 minutes in 100 ul MEM media supplemented with 0.3% BSA and trypsin-TPCK (1ug/m). The cells were washed twice with PBS and the virus-serum mixture was transferred onto the cells. At 18 h post infection, cells were washed once and 30ul of 0.5% NP-40 in DPBS added for 15 minutes at 37°C. Plates were read using Synergy H1 at ex 580 / em 610nm. The average signal measured in absence of the Ab was considered 100% infectivity and used for calculation of fraction infected at any given Ab concentration.

#### Infected cell binding assay

MDCK cells were infected using PR8-mCherry expressing virus at MOI=5 for 5h at 37°C. After incubation, cells were transferred into tubes and stained with pre-immune or immune sera diluted 1:50 for 90 min at 37°C. After washes with PBS/0.1% BSA cells were fixed overnight in 1.5% PFA. Cells were washed and stained with Pe-Cy7 anti-mouse IgG (poly4053)(Biolegend, cat405315) for 30 min at 4°C before acquisition via flow cytometry.

#### Statistical analysis

GraphPad Prism (GraphPad Software Inc.) was used for statistical analysis. Mice were randomly allocated to experimental groups and the investigators were not blinded on the identity of the samples except for the neutralization and ADCS and ADCP assays. For comparison between two groups unpaired student’s t test was performed. For comparison of one variable between multiple groups one-way ANOVA with Tukey’s multiple comparison test was performed. For comparison between multiple variable across multiple groups two-way ANOVA with Holm-Sidak’s multiple comparison test was performed. When data did not pass the normality test it was log transformed before plotting and statistical analysis. For all figures data point indicate individual mice. * represents P < 0.05, ** P < 0.01, *** P < 0.001, **** P < 0.0001.

---

**Figure S1.**
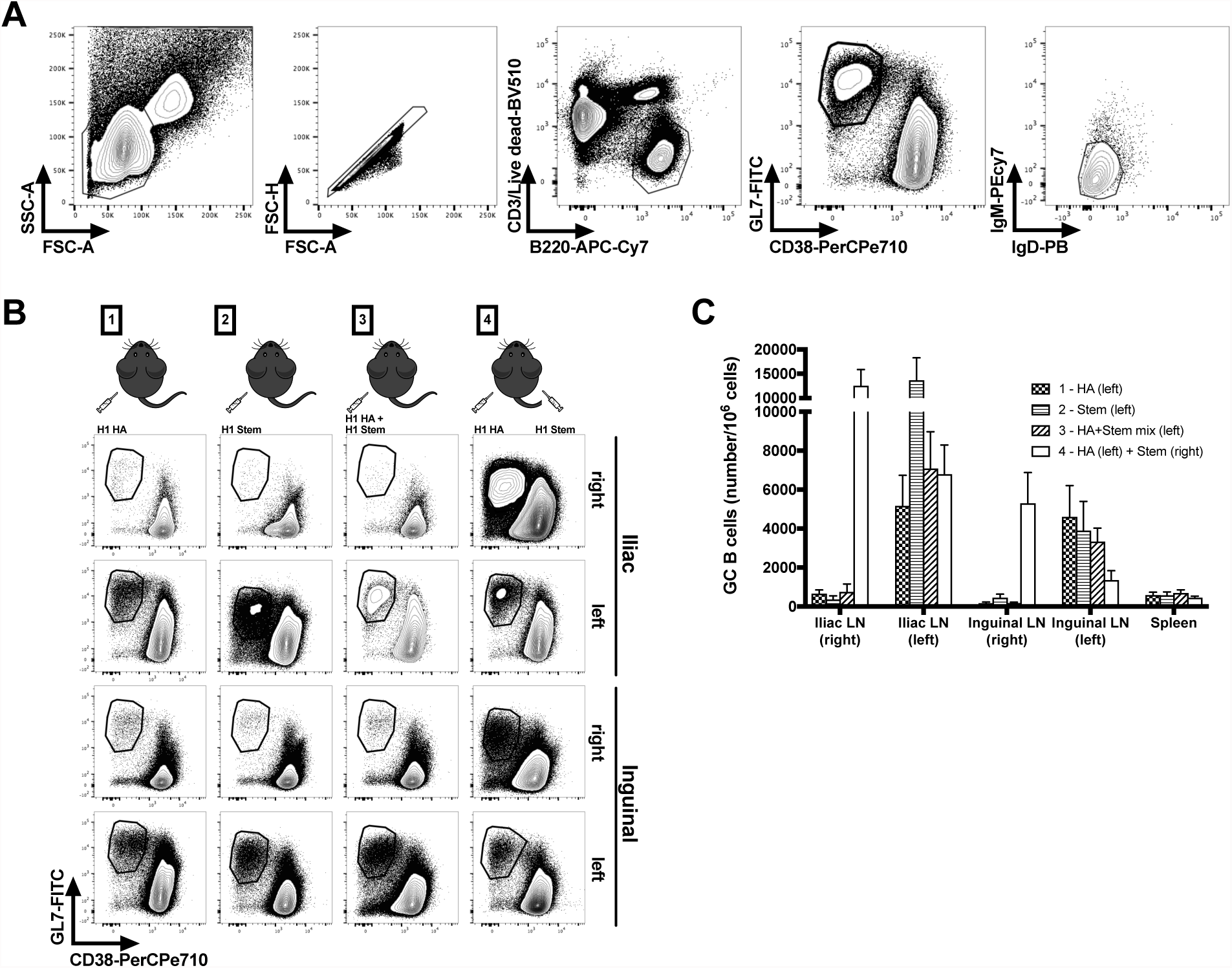
Flow cytometry gating strategy and GC B cell quantification. (A) gating strategy for flow cytometry, cells were gated as live CD3^-^ B220^+^ GL7^+^ CD38^-^ IgD^-^ IgM^-^ (B) Representative flow cytometry plot showing size of GC B cells, gated as live CD3^-^ B220^+^ GL7^+^ CD38^-^ (C) Frequency of GC B cells as detected in the different LNs and spleen after immunization as in Fig. 1A (n=9). Three independent experiments with 4 mice each (pooled for the first experiment). Bar graph represent mean and bars SEM

**Figure S2.**
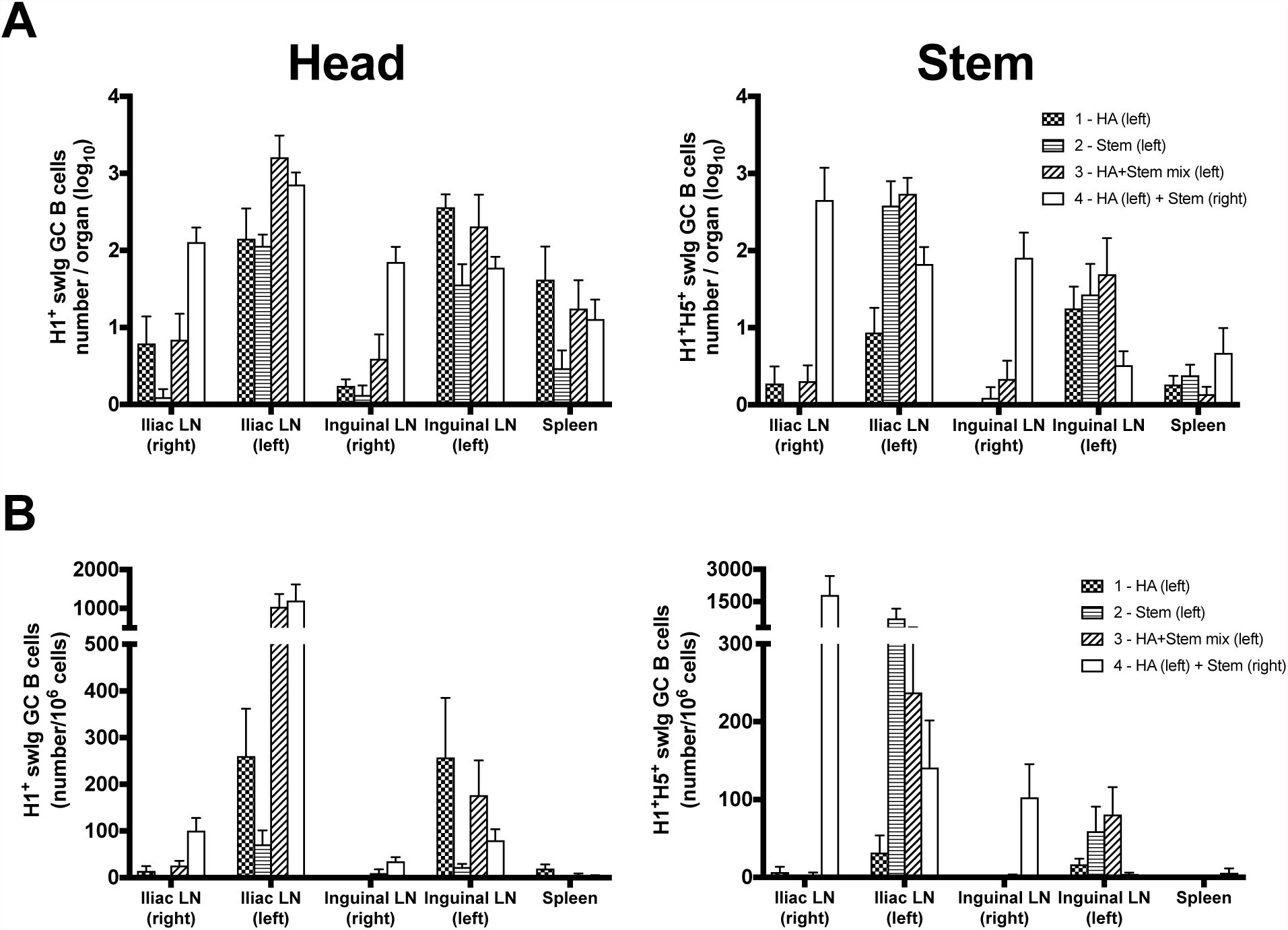
Distribution of head and stem specific GC B cells in lymphoid organs. Absolute number (A) and frequency (B) of swig GC B cells as detected in the different LNs and spleen after immunization (n=9) as in Fig 1C. Three independent experiments with 4 mice each (pooled for the first experiment). Bar graph represent mean and bars SEM

**Figure S3.**
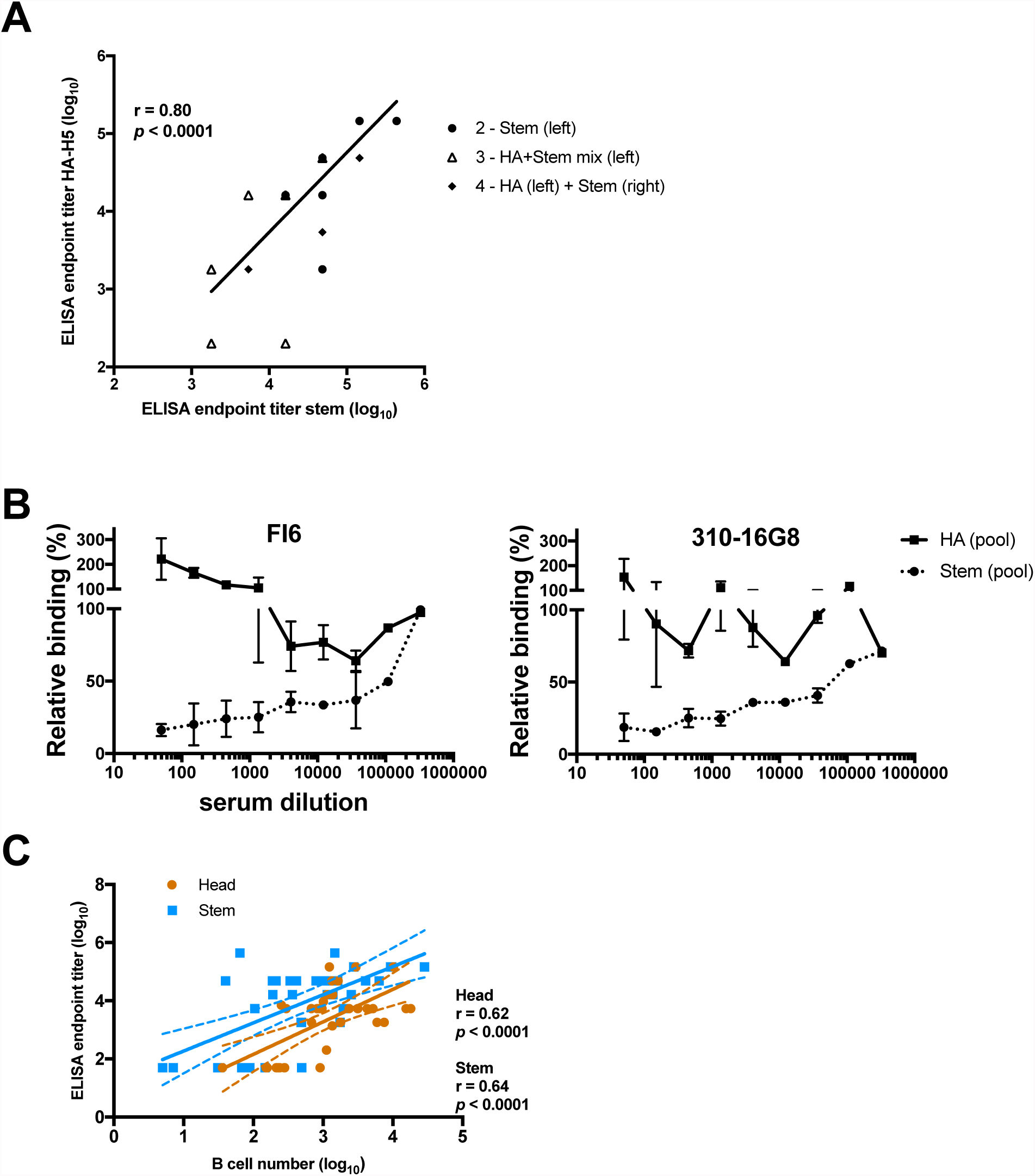
Characterization of serum Abs. (A) Correlation showing endpoint titers of sera on stem-only antigen (as in Fig. 2A) vs H5 antigen for mice with a stem titer (groups 2, 3 and 4) (n=34). (B) Competition of stem mAbs FI6 and 16G8 with pooled stem sera (from group 2 immunization) or HA sera (from group 1) for binding on HA-PR8. PR8-HA coated plates were incubated with serial dilution of pooled sera and subsequently detected with stem mAbs at EC_75_. Two independent experiments with 12 pooled sera each. Bars is SEM. (C) Correlation of Head/Stem specific swig GC B cells as in Fig. 1C vs ELISA endpoint titers as in Fig. 2A

**Figure S4.**
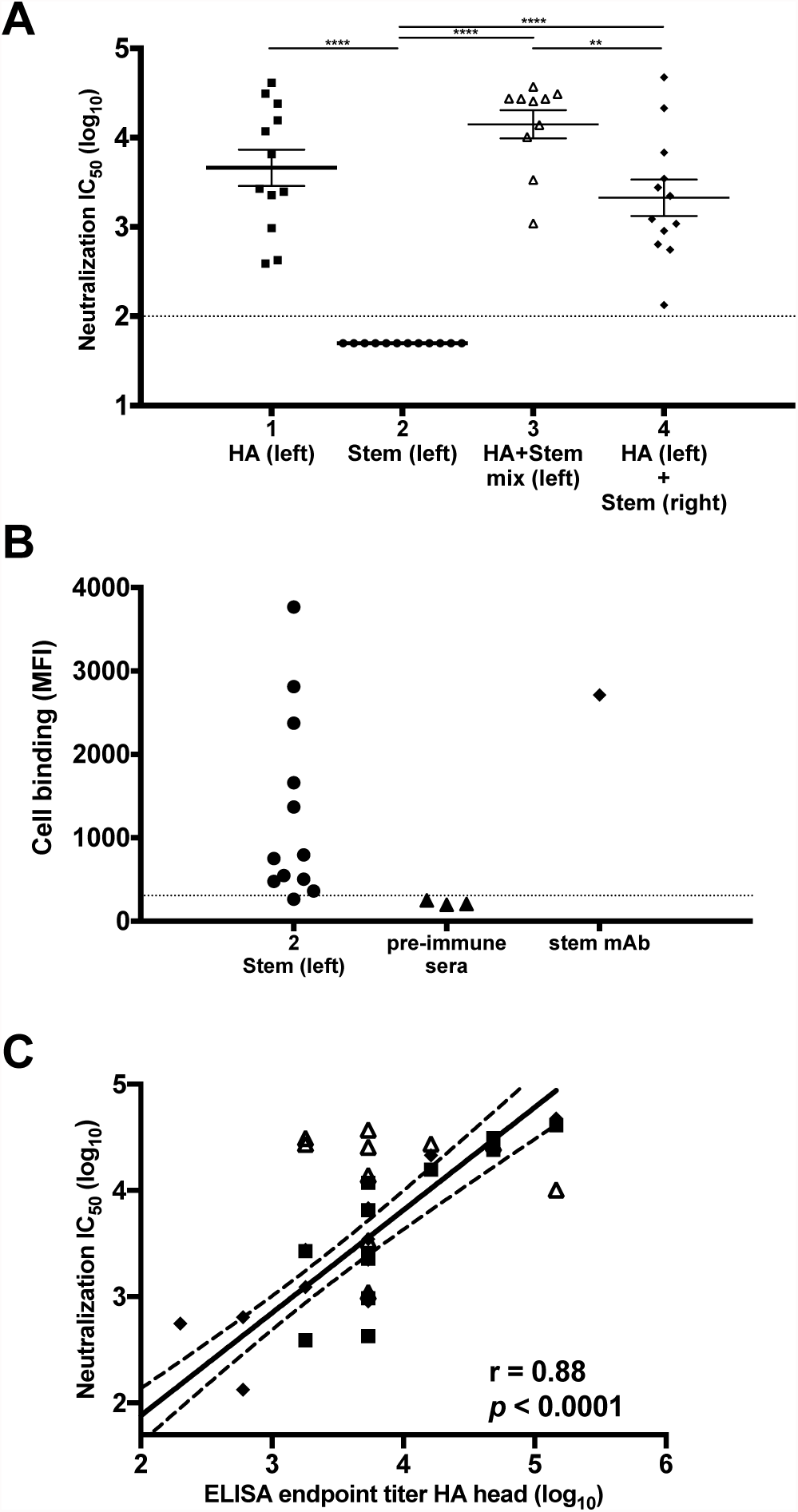
Neutralization and cell binding capacity of sera. (A) Sera from the different groups (n=12) was tested for its ability of neutralizing PR8 virus. Results are expressed as IC_50_, the serum dilution that gave half of the maximal inhibition. Three independent experiments with 4 mice each (n=12 for groups 1, 2, 4 and n=10 for group 3) and 4 technical replicates for each serum. Bar is mean ± SEM, statistical analysis was performed using one-way ANOVA with Tukey multiple comparison test. (B) Sera from animals immunized with stem only (group 2) (n=12) was tested for its ability to bind PR8-infected cells. Dotted line indicates cut-off given by the average of 3 pre-immune sera+3SD. Stem mAb FI6 was also tested as control. (C) correlation between neutralization IC_50_ as in A and ELISA endpoint titer for HA head as in Fig. 2A

**Fig. S5.**
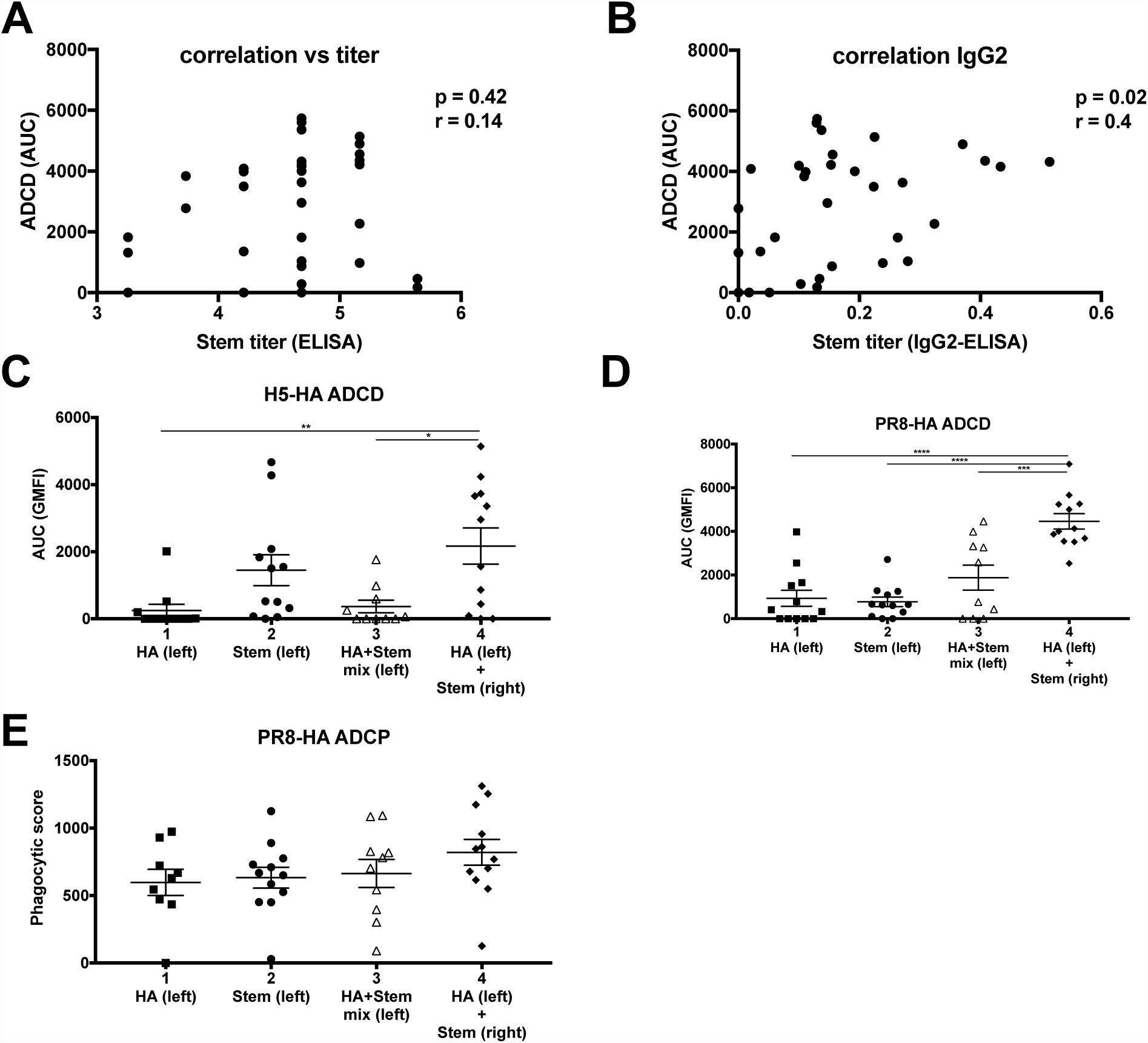
Effector function activity of sera. Correlation of ADCD response as in Fig. 2C with total stem Abs titer (as in Fig. 2A) (A) or stem-specific IgG2 (As in Fig. 2B)(B). Sera were tested for the ability to induce ADCD on H5-HA- (C) or PR8-HA (D) conjugated beads. Data is presented as area under the curve (AUC) of geometrical mean fluorescent intensity (GMFI) of 1:5 and 1:10 dilutions, and is the mean of two technical replicates (n=12 for groups 1, 2, 4 and n=10 for group 3).</br>(E) Ability of the sera to induce ADCP on PR8-HA-conjugated beads by primary monocytes. Each data point is the mean of two technical replicates (n=12 for groups 1, 2, 4 and n=10 for group 3). Three independent experiments with 4 mice each. Bar is mean ± SEM, statistical analysis was performed using one-way ANOVA with Tukey multiple comparison test.

**Fig. S6.**
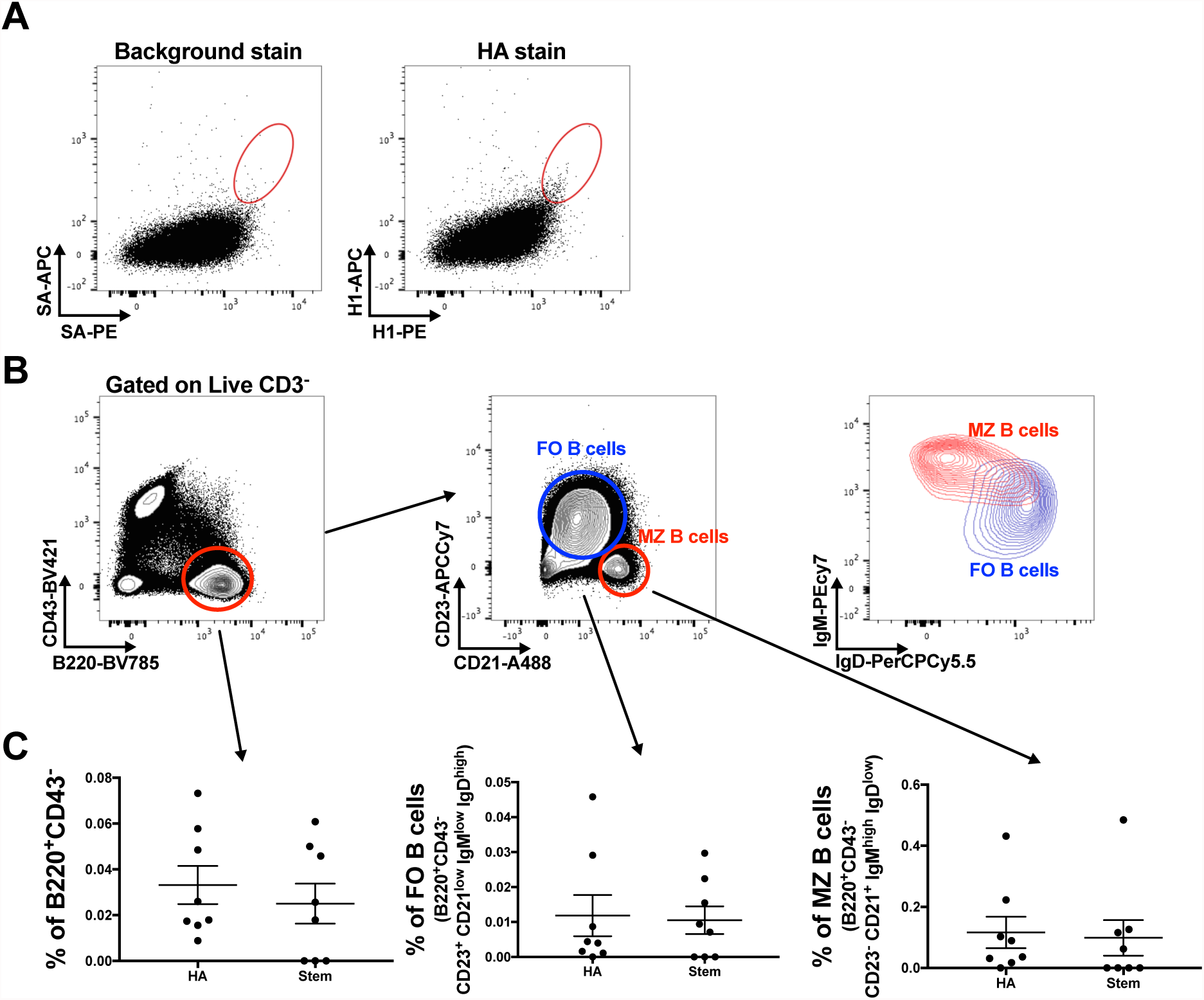
Naive B cell precursor frequency is similar between head and stem. (A) Representative flow cytometry plot showing background stain of naïve B cells (CD3^-^ CD43^-^ B220^+^) on streptavidin (SA) conjugated to PE and APC and used in combination and staining of the same cells with PR8-HA. (B) Gating strategy for total mature naïve B cell population (CD3^-^ CD43^-^ B220^+^), follicular (FO) (CD3^-^ CD43^-^ B220^+^ CD23^+^ CD21^low^ IgM^low^ IgD^high^) and marginal zone (MZ) (CD3^-^ CD43^-^ B220^+^ CD23^-^ CD21^l+^ IgM^high^ IgD^low^) B cells. Overlay shows IgM and IgD expression in FO vs MZ B cells. (C) Precursor frequency as % of parent population for mature naïve, FO and MZ B cells. (n=8). Three independent experiments with 2 or 4 mice each. Bar graph represent mean and bars SEM, statistical analysis was performed using unpaired t-test

**Fig. S7.**
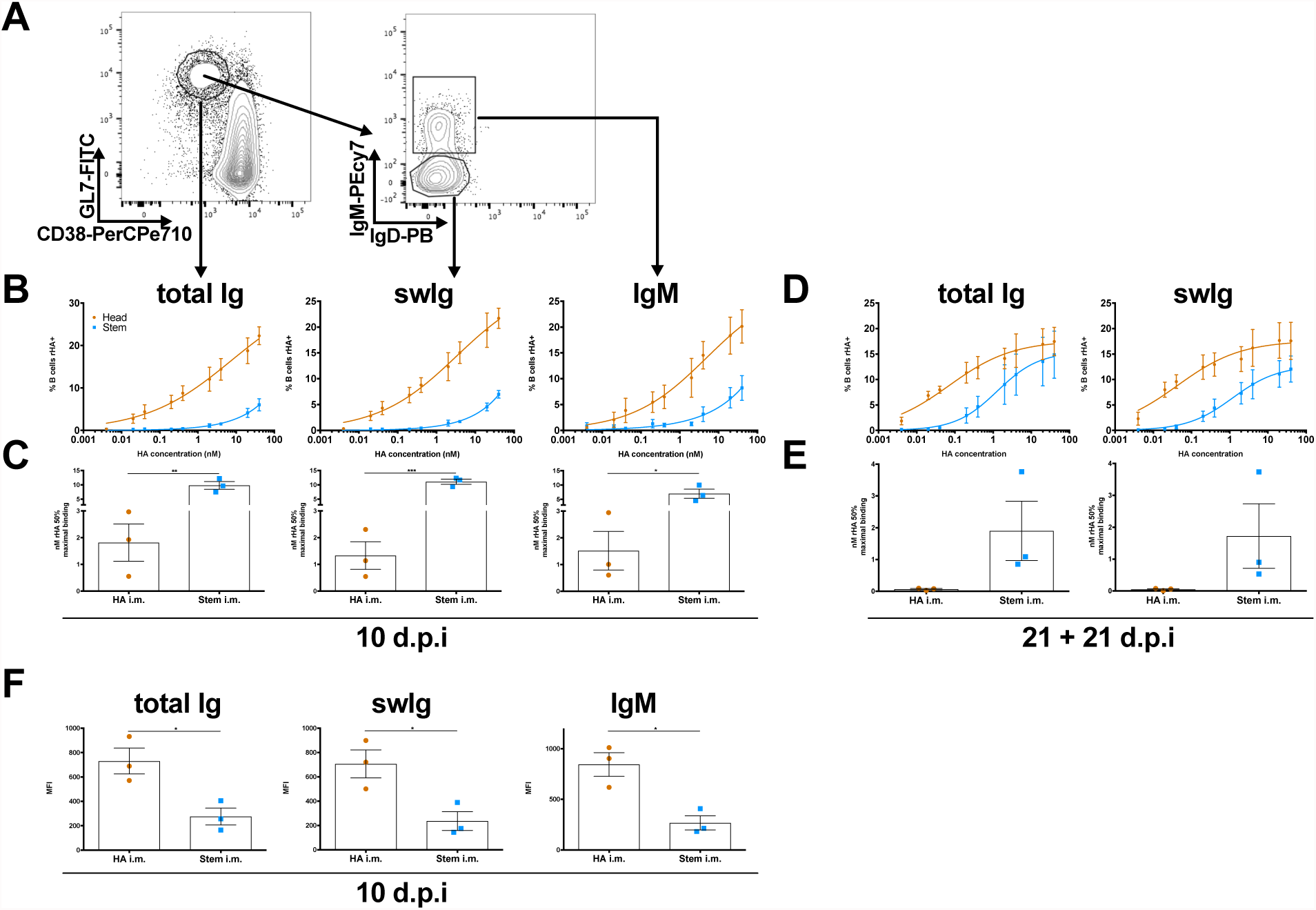
Early B cell affinity is different for head *vs* stem. (A) Representative flow cytometry plot showing total (CD3^-^ B220^+^ GL7^+^ CD38^-^), swig (CD3^-^ B220^+^ GL7^+^ CD38^-^ IgD^-^ IgM^-^) and IgM (CD3^-^ B220^+^ GL7^+^ CD38^-^ IgD^-^ IgM^+^) GC B cells and the gating selection for the AC_50_ at 10 days post HA or stem i.m. Pooled iliac LNs from 3 mice at 10 days after immunization (B) or at 21 days after challenge (D) with full length PR8 HA (orange) or stem (blue) were stained with graded amount of HA-PR8 (orange) or HA-H5 (blue) and plotted against frequency of GC B cells stained (n=3). (C-E) nM concentration of HA giving half maximal binding (AC_50_) was derived from individual experimental curves (n=3). (F) Median fluorescence intensity (MFI) of HA^+^ B cells at the highest concentration (66nM) at 10 d.p.i. Three independent experiments with 3 pooled LN each. Bar graph represent mean and bars SEM, statistical analysis was performed using unpaired t-test

**Figure S8.**
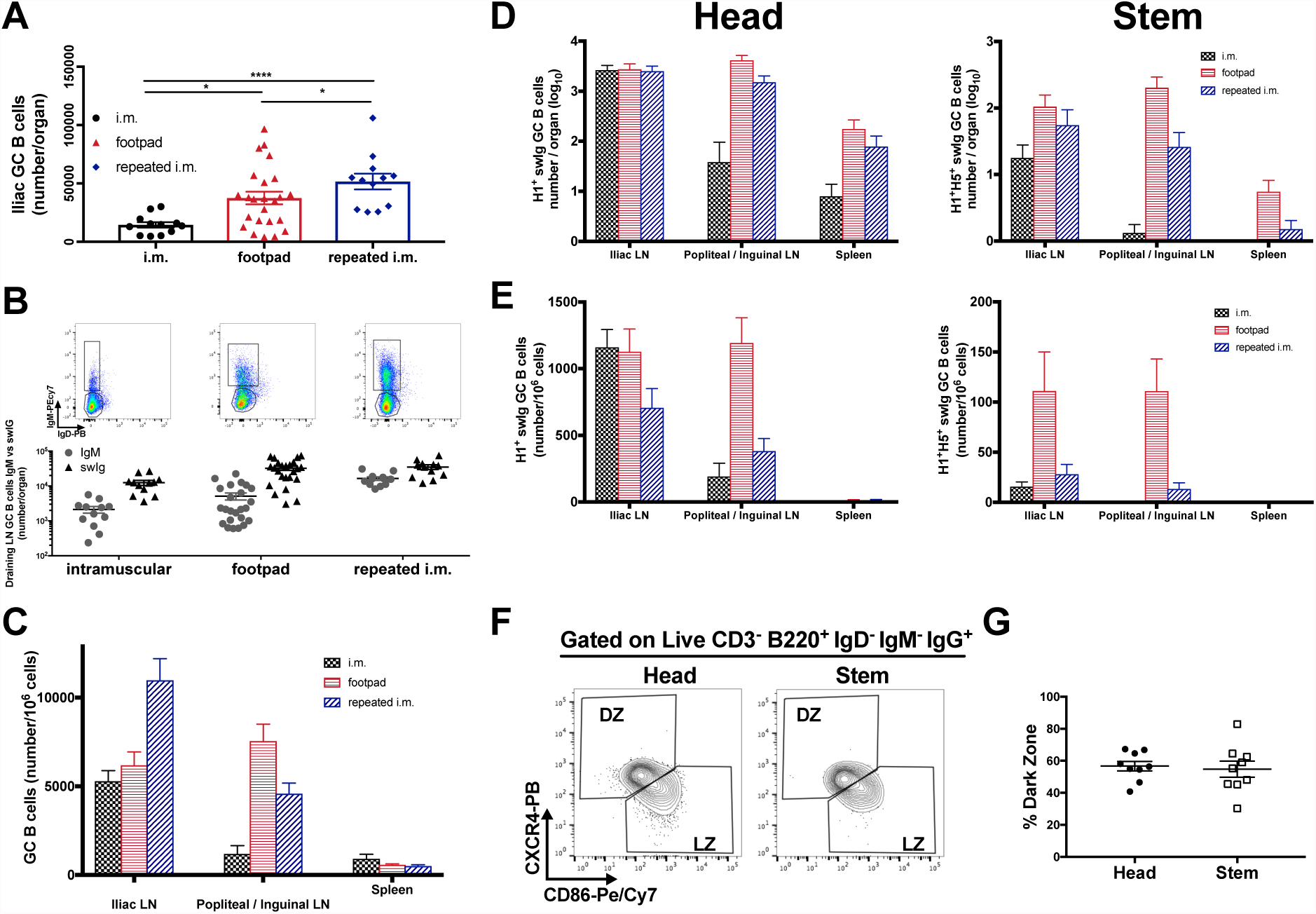
GC characterization upon i.m., f.p. or repeated i.m. immunizations. (A) Number of total Iliac GC B cells after immunization. (B) Representative flow cytometry plot showing draining LN GC B cells (gated as live CD3^-^ B220^+^ GL7^+^ CD38^-^ IgD^-^) and further dividing them into swig (IgM^-^) or IgM^+^. (C) Frequency of GC B cells depending on draining LN (popliteal for f.p. and inguinal for i.m. and repeated i.m.). Absolute number (D) and frequency (E) of swig GC B cells as detected in the different LNs and spleen after immunization as in Fig. 3C. (n=12 for i.m. and repeated i.m. and n=24 for f.p.). Three independent experiments with 4 mice each for i.m and repeated i.m and 5 independent experiments with 5 or 4 mice each for f.p. Bar is mean ± SEM, statistical analysis was performed using one-way ANOVA with Tukey multiple comparison test.</br>(F) Representative flow cytometry plot showing Dark zone (DZ) (CD3^-^ B220^+^ GL7^+^ CD38^-^ IgD^-^ IgM^-^ IgG^+^ CXCR4^+^ CD86^-^) *vs* light zone (LZ) (CD3^-^ B220^+^ GL7^+^ CD38^-^ IgD^-^ IgM^-^ IgG^+^ CXCR4^-^ CD86^+^) distribution of Head or stem specific GC B cells at 11 after f.p. immunization. (G) Quantification of the frequency of Head or stem GC B cells in the DZ (n=9). Two independent experiments with 5 and 4 mice each. Bar graph represent mean and bars SEM, statistical analysis was performed using unpaired t-test

**Figure S9.**
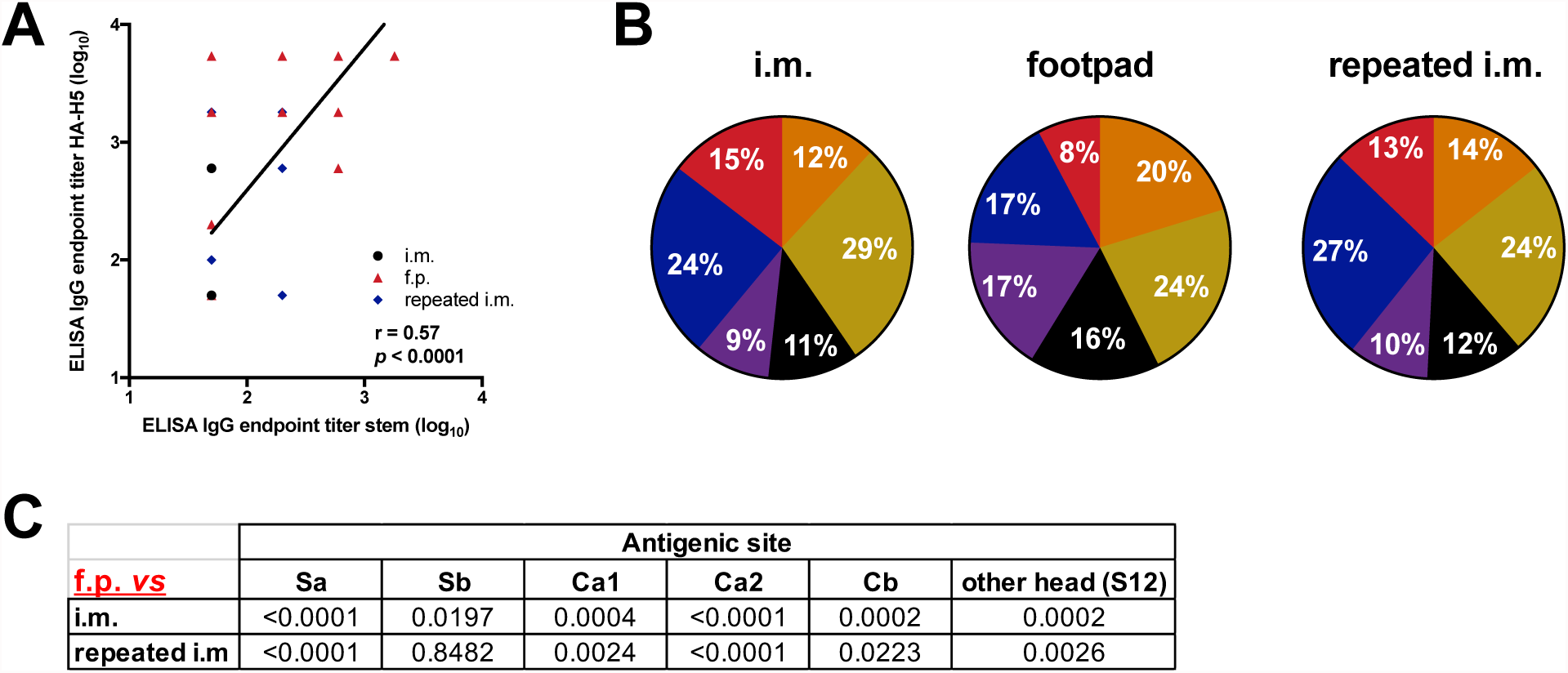
Characterization fo serum Abs. (A) Correlation showing endpoint titers of sera on stem-only antigen (as in Fig. 3F) vs H5 antigen (n=12 for i.m. and repeated i.m. and n=24 for f.p.). (B) Serially diluted sera from i.m., f.p. or repeated i.m. were tested by ELISA on a panel of HA from drifted PR8 viruses to determine their immunodominance profile. Shown is a pie chart graph showing the AUC for each of the canonical antigenic sites (Sa= blue, Sb= gold, Ca1= purple, Ca2= orange, Cb= red, non-canonical head site= black). (C) Statistical analysis related to B comparing f.p. with the other two immunization regimes. The analysis was performed using one-way ANOVA with Tukey multiple comparison test for each of the antigenic sites. No difference was detected between i.m. and repeated i.m. for any of the sites. (n=12 for i.m. and repeated i.m. and n=23 for f.p.).

**Fig. S10.**
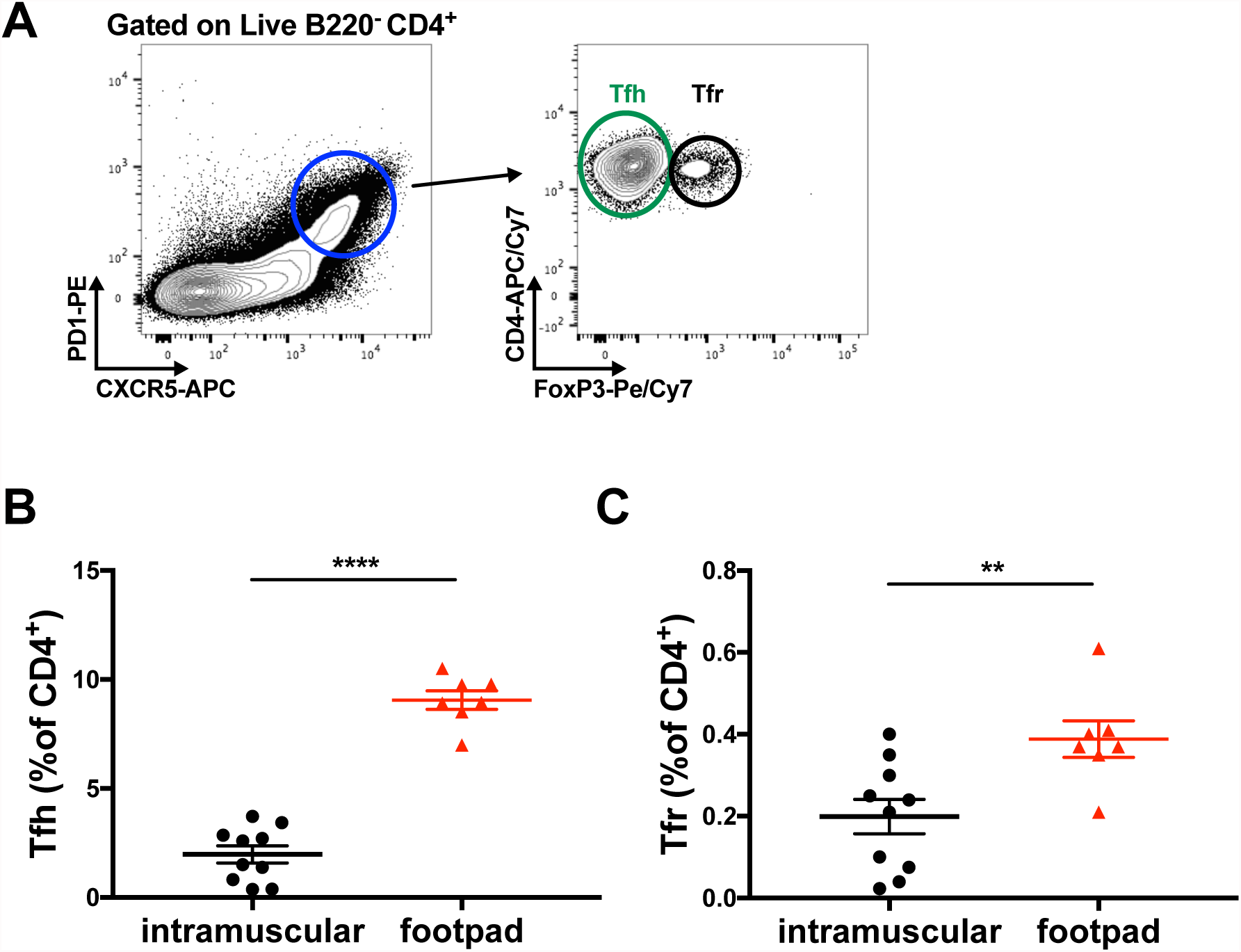
F.p. immunization alters T follicular cells frequency. (A) Gating strategy for T follicular (Tf) cell population (B220^-^ CD4^+^ CXCR5^+^ PD1^+^) and further classification into T follicular helper (Tfh) and T follicular regulatory (Tfr) based on FoxP3 expression. Tfh (B) and Tfr (C) expressed as frequency of CD4^+^ T cells (n= 10 for i.m. and 7 for f.p.). Two independent experiments with 5 mice each for i.m. and with 4 and 3 mice for f.p. Bar is mean ± SEM, statistical analysis was performed using unpaired t-test

